# Estimating the proportion of bystander selection for antibiotic resistance among potentially pathogenic bacterial flora

**DOI:** 10.1101/288704

**Authors:** Christine Tedijanto, Scott W. Olesen, Yonatan H. Grad, Marc Lipsitch

## Abstract

Bystander selection ‐ the selective pressure for resistance exerted by antibiotics on microbes that are not the target pathogen of treatment ‐ is critical to understanding the total impact of broad-spectrum antibiotic use on pathogenic bacterial species that are often carried asymptomatically. However, to our knowledge, this effect has never been quantified. We quantify bystander selection for resistance for a range of clinically relevant antibiotic-species pairs as the proportion of all antibiotic exposures received by a species for conditions in which that species was not the causative pathogen (“proportion of bystander exposures”). Data sources include the 2010-2011 National Ambulatory Medical Care Survey and National Hospital Ambulatory Medical Care Survey (NAMCS/NHAMCS), the Human Microbiome Project, and additional carriage and etiological data from existing literature. For outpatient prescribing in the United States, we find that this proportion over all included antibiotic classes is over 80% for 8 of 9 organisms of interest. Low proportions of bystander exposure are often associated with infrequent bacterial carriage or concentrated prescribing of a particular antibiotic for conditions caused by the species of interest. Applying our results, we roughly estimate that pneumococcal conjugate vaccination programs result in nearly the same proportional reduction in total antibiotic exposures of *S. pneumoniae*, *S. aureus*, and *E. coli*, despite the latter two organisms not being targeted by the vaccine. These results underscore the importance of considering antibiotic exposures of bystanders, in addition to the target pathogen, in measuring the impact of antibiotic resistance interventions.

**Significance Statement:** Bystander selection, defined as the inadvertent pressure imposed by antibiotic treatment on microbes other than the targeted pathogen, is hypothesized to be a major factor in the propagation of antibiotic resistance, but its extent has not been characterized. We estimate the proportion of bystander exposures across a range of antibiotics and potential pathogens commonly found in the normal flora and describe factors driving variability of these proportions. Impact estimates for antibiotic resistance interventions, including vaccination, are often limited to effects on a target pathogen. However, the reduction of antibiotic treatment for conditions caused by one pathogen may have the broader potential to decrease bystander selection pressures for resistance on many other organisms.

## Introduction

Antibiotic use creates selective pressures favoring resistant microbes. While designed to control the pathogenic bacteria causing an infection (we use the term “target pathogen”), currently available antibiotics often have antimicrobial activity against many bacterial species and disseminate widely throughout the body (1). Thus, the bacteria that comprise the human microbiome are subject to the selective pressures applied by most antibiotic consumption (2–4). These selective pressures for resistance experienced by microbial flora due to antibiotic exposures for a condition not caused by that species can be called “bystander selection”. While non-pathogenic commensals residing in the microbiome are always bystanders, opportunistic and obligate pathogens, which are often carried asymptomatically, lie at the critical intersection where resistance is clinically relevant and the extent of bystander selection among them is unknown. Although commensals may be an important reservoir for resistance elements, our ability to measure horizontal transmission between organisms in the microbiome is limited. Therefore, we focus on bystander selection for resistance due to antibiotic exposures experienced directly by potential pathogens.

Quantifying bystander selection, as defined above, is important for evaluating the impact of antibiotic resistance control interventions. For example, vaccination and infection control strategies are designed to reduce the need for appropriate antibiotic treatments and thus decrease selective pressure for resistance on a target pathogen: vaccination by reducing the incidence of disease from, say, *Streptococcus pneumoniae* and thus the need for antibiotic treatment (5, 6), and infection control by reducing the incidence of hospital-acquired infections that will require treatment. Often overlooked is the impact that averted treatment may have beyond the target pathogens, because each treatment averted would have exerted selection on bystanders as well. For stewardship interventions, which aim to avert inappropriate treatment of conditions that are never or seldom caused by bacteria, the primary goal of the intervention is to avert bystander selection of the patient’s normal flora. Mathematical transmission models that aim to simulate the dynamics of antibiotic resistance and to project the impact of interventions on pathogenic bacteria with an asymptomatic carriage state often assume that treatment incidence is independent of colonization with the bacterium of interest, implying that bystander selection is the rule rather than the exception (7–9). Prior to this study, there has not been sufficient evidence to support this claim.

This work aims to estimate the extent of bystander selection for resistance due to outpatient prescribing in the US for a range of clinically relevant species and antibiotic combinations. Prescriptions are used as a measured proxy for exposures and, ultimately, for selection. We use existing data, including the National Ambulatory Medical Care Survey and National Hospital Ambulatory Medical Care Survey (NAMCS/NHAMCS) to estimate prescription volume and associated diagnoses, and the Human Microbiome Project and other studies of bacterial carriage to estimate the microbial communities subject to selection. *We quantify bystander selection as the proportion of total exposures of an antibiotic experienced by a species when that species was not the target pathogen of treatment, and will refer to this measure as the “proportion of bystander exposures”*.

Understanding the contribution of bystander exposures to the landscape of selective pressures for antibiotic resistance at the population level will help to inform interventions including vaccines and antibiotic stewardship. Given the special attention of the current issue of *PNAS* to vaccines and antimicrobial resistance, we spell out how such measures can contribute to estimating the impact of vaccines, in particular pneumococcal conjugate vaccines, whose impact on antimicrobial resistance has received arguably the most attention of any vaccine (10, 11).

## Results

### Data source characteristics

After exclusion of visits resulting in hospital or observation unit admission, the National Ambulatory Medical Care Survey (NAMCS) and National Hospital Ambulatory Medical Care Survey (NHAMCS) from 2010-2011 with nationally representative sampling weights were used to estimate outpatient diagnosis and prescription volume in the US (see Materials and Methods). 94.4% of total sampled visits were included in the analysis, with 14.7% of these visits resulting in at least one antibiotic prescription. Visits with one or more of the conditions explicitly included in the bystander analysis (x-axis of Figure 1A) accounted for 55.8% of unweighted prescriptions of our antibiotic classes of interest. Of included visits with one of these conditions, 6% were clinical encounters with patients less than one year old and 19% with patients between the ages of 1 and 5. The Human Microbiome Project (HMP) data includes isolates from healthy individuals between the ages of 18 and 40 sampled at 15-18 locations across five major body sites: the nasal passages, oral cavity, skin, gastrointestinal tract, and urogenital tract. This analysis was agnostic to body site; an individual was considered to have positive carriage status of a particular species if that species was identified at any body site.

**Figure 1.**
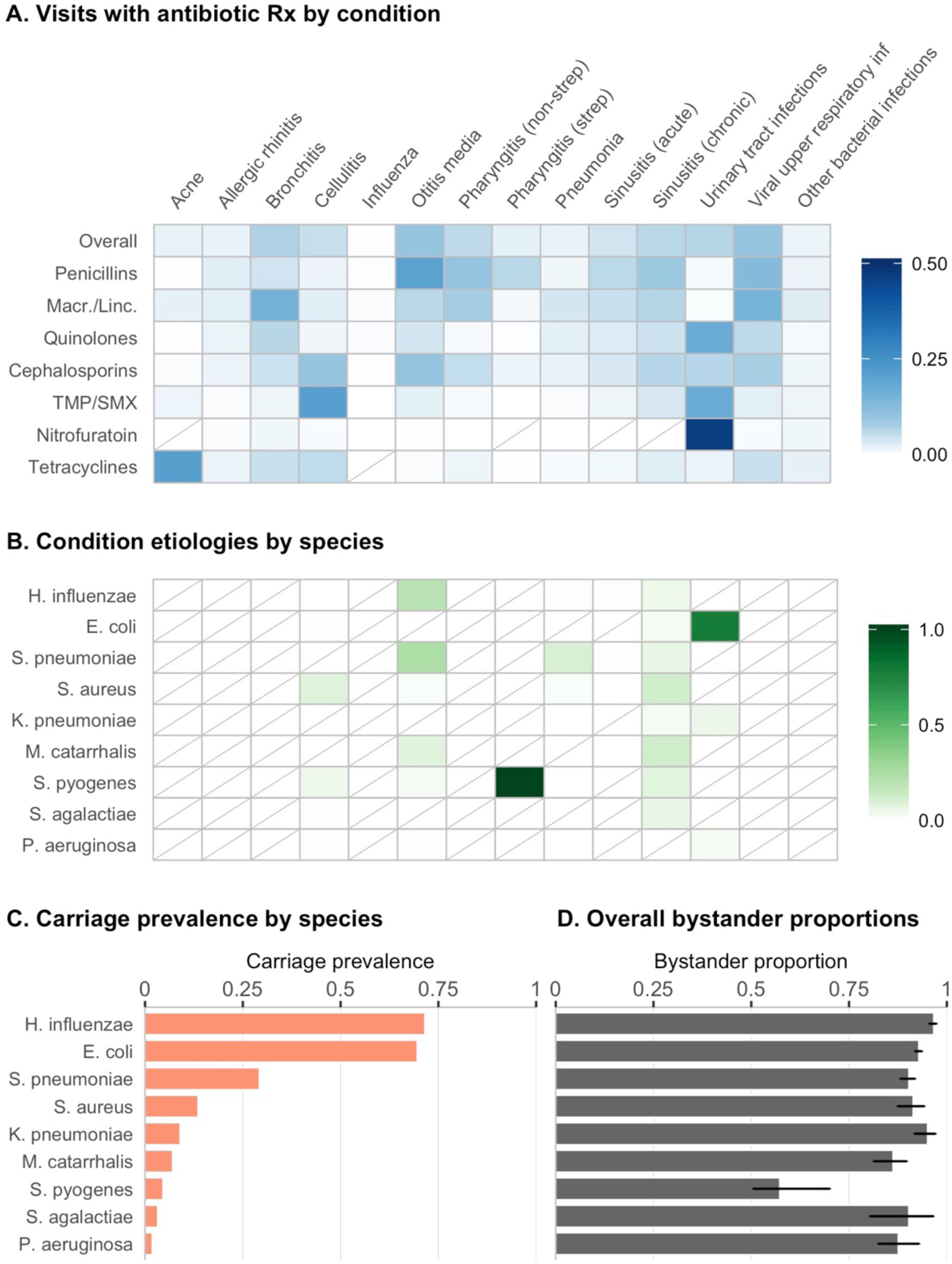
Inputs and overall results of bystander analysis. ***A:*** Heat map shading represents the proportion of visits (after weighting to be nationally representative) with a diagnosis of the specified condition, given that the visit resulted in a prescription of the specified antibiotic class. Results for TMP/SMX and nitrofurantoin are for the individual drug instead of an antibiotic class. Rows are not required to sum to 100%, as only a subset of conditions are shown, and each visit may be associated with more than one condition. Antibiotics included in each class are based on the Multum Lexicon classification system. Macr./Linc. = Macrolides/lincosamides; TMP/SMX = trimethoprim-sulfamethoxazole. Diagonal lines indicate cells with value of 0. ***B:*** Heat map shading represents the estimated etiology of each condition by species. If etiological data was available for multiple age groups, the weighted mean based on the relative frequency of visits (after weighting to be nationally representative) for that condition is shown. Diagonal lines indicate cells with value of 0. ***C:*** Bars indicate mean carriage prevalence of each species across age groups, weighted by relative frequency of visits (after weighting to be nationally representative). ***D:*** Bars indicate proportion of bystander exposures by antibiotic class and species >(*B*_*as*_) with 95% confidence intervals. “Overall” estimates reflect exposures to antibiotic in any of the classes shown in Panel A.

### Estimating the proportion of bystander selection for resistance, by species and antibiotic or class

The three inputs required to calculate the proportion of bystander exposures are antibiotic prescriptions by condition, condition etiologies, and carriage prevalence of each species. In Figure 1A, we depict antibiotic use as the proportion of weighted visits at which the specified condition was diagnosed, given that the visit resulted in a prescription of the specified antibiotic class; this value is a function of both antibiotic use by condition and volume of visits for the given condition. For example, this proportion is relatively high for use of penicillins for suppurative otitis media and macrolide/lincosamide use for bronchitis, common conditions leading to frequent prescriptions of the respective antibiotic class. While a high proportion of pneumonia cases also result in antibiotic prescriptions, low incidence in the outpatient setting leads to low proportions of antibiotic use associated with this condition. Nitrofurantoin presents an extreme case where use is targeted towards a single, common condition, urinary tract infections (UTIs). While many of the conditions that we consider do not have bacterial etiologies, those that do are infrequently caused by our bacteria of interest; conditions which are primarily caused by a single organism are streptococcal sore throat (strep), which we assume is always caused by *S. pyogenes*, and UTIs, commonly caused by *E. coli* (Figure 1B). Carriage prevalence varies dramatically from nearly 75% for *H. influenzae* and *E. coli* to well below 5% for *S. pyogenes, S. agalactiae*, and *P. aeruginosa* (Figure 1C).

The connections between antibiotic use, etiology, and carriage prevalence may be observed more clearly when considering bystander selection for a single antibiotic or class. We discuss two examples ‐ *E. coli* and quinolones, and *S. pyogenes* and penicillins. Quinolones, such as ciprofloxacin, are frequently used to treat UTIs (Figure 1A), which, as previously mentioned, are commonly caused by *E. coli* (Figure 1B). This alignment between antibiotic use and etiology results in a lower proportion of bystander exposures to quinolones for *E. coli*, compared to other species (SI Appendix, Figure S1). This relationship is accentuated for species with low carriage prevalence. For example, a moderate proportion of penicillin use is directed towards strep throat (Figure 1A), for which we assume *S. pyogenes* is the sole cause (Figure 1B). Due to this extreme etiology and low carriage prevalence, exposures of *S. pyogenes* to all included antibiotics, especially penicillins, occur more frequently when *S. pyogenes* is a target pathogen and not a bystander. This factor also contributes to the low bystander proportion of *P. aeruginosa* for antibiotics used to treat UTIs. The bystander proportion for *P. aeruginosa* is often comparable to that of *E. coli*, even though it is a far less common cause of UTIs; compare this to *K. pneumoniae*, which causes more UTIs than *P. aeruginosa*, but is more prevalent in carriage and thus experiences more bystander exposures. Thus, low carriage prevalence is also a driver of low bystander selection.

Overall, the proportion of bystander exposures exceeded 80% for 8 out of 9 organisms (all except *S. pyogenes)* when considering exposures to any of our antibiotic classes of interest (Figure 1D), and was above 80% for 133 out of 153 (86.9%) antibiotic-species pairs (SI Appendix, Figure S4). These results indicate that for the majority of antibiotic and species combinations, fewer than 20% of the exposures of that species to the antibiotic in question occur in the context of treating a disease caused by that species. Of particular clinical interest, 83.9% (95% CI: 80.9%, 86.6%) of *S. pneumoniae* exposures to penicillins and 93% (95% CI: 90.7%, 94.5%) of exposures to macrolides occurred when *S. pneumoniae* was not the target pathogen of disease. For *E. coli*, the proportion of bystander exposures was 81.3% (95% CI: 79.1%, 84%) for quinolones and 93.2% (95% CI: 91.4%, 94.3%) for cephalosporins, both of which are often used to treat pathogenic *E. coli*. The proportion of bystander exposures was similarly high for *S. aureus* and penicillins at 91% (95% CI: 86%, 94.6%). *S. pyogenes*, which is rarely carried asymptomatically, only experienced 35.2% (95% CI: 30.6%, 55.9%) of its exposures to penicillins as a bystander.

Antibiotic resistance in *N. gonorrhoeae* is of urgent concern, and recent ecological (12) and individual-level (13) studies have implicated bystander selection as a potential driver of macrolide resistance. Due to the low incidence of gonorrhea in the general population, limited data was available from NAMCS/NHAMCS. We used additional data from the Gonococcal Isolate Surveillance Project (GISP) (14) with slightly modified methods (see Materials and Methods) to estimate the proportion of bystander exposures for *N. gonorrhoeae*. From 2010-2011, the proportion of bystander exposures for *N. gonorrhoeae* was 97.7% for ciprofloxacin and 4.8% for ceftriaxone. At the antibiotic class level, the proportion of bystander exposures for *N. gonorrhoeae* was 97.5% for quinolones and 14.6% for cephalosporins. GISP data on macrolide and tetracycline use were unavailable for 2010. For 2011, GISP reports combine prescription data for azithromycin and erythromycin and for doxycycline and tetracycline. Assuming that 75-95% of the azithromycin-erythromycin volume can be attributed to azithromycin, we estimate that the proportion of bystander exposures for *N. gonorrhoeae* ranges from 22.8% to 18.9%, respectively. Similarly, assuming that 75-95% of reported doxycycline-tetracycline use is doxycycline results in estimates for the proportion of doxycycline bystander exposures for *N. gonorrhoeae* of 29.7% to 25%. At the antibiotic class level, the proportion of bystander exposures for *N. gonorrhoeae* was 22.8% for macrolides/lincosamides and 28.6% for tetracyclines, accounting for all doxycycline-tetracycline use.

### Application to vaccine impact on antimicrobial exposures in bystanders

We provide here preliminary estimates of the bystander impact of vaccines to illustrate how such calculations might be performed. Because the requisite quantities have not all been estimated in the same population, we combine estimates from different populations for the purposes of illustration, but we note the need for additional data to improve the level of confidence in such calculations by comparing quantities within a single population. We take the example of the pneumococcal conjugate vaccine (PCV), which reduces bacteremia, meningitis, pneumonia and otitis media caused by 7, 10 or 13 serotypes of pneumococci, depending on the formulation.

Considering only immediate direct effects, we made a conservative estimation of the effect of PCV on antimicrobial exposures of bystander organisms (see Materials and Methods). A randomized controlled trial of the seven-valent pneumococcal conjugate vaccine found a 7.8% reduction in otitis media among vaccinated vs. control children, and a 5.4% in all-cause antimicrobial prescribing, mainly attributed to reduced otitis media (5). Restricting attention to the 0-1 and 1-5 year old age groups in our study, this would translate to a 5.2% reduction in exposure of *S. aureus* to antibiotics, and a 5.4% reduction in exposure of *E. coli* to antibiotics in these age groups.

Much larger estimates of impact on otitis media are obtained in studies that account for herd immunity effects and for the possibility that PCVs can indirectly prevent some non-pneumococcal otitis media (15). An Israeli study found a 57-71% reduction in all-cause otitis media associated with the rollout of PCV13 in various age groups up to the third birthday (16), while a study in the UK found a 36% reduction in otitis media among children under 10 years old comparing the post-PCV13 period to the pre-PCV7 period, and a 29% reduction in otitis media-associated antimicrobial prescribing for the same comparison (6). Impact on total antimicrobial prescribing was not reported in the UK (6). If we assume that the ratio of 0.69 percentage points reduction in total prescribing per percentage point reduction in otitis media can be extrapolated from the RCT in California (5), this 36% reduction in otitis media would correspond to a 25% reduction in all-cause antibiotic prescribing. Using our estimates of prevalence and bystander proportion, this would yield a 24.99% and 24.2% reduction in outpatient exposure of bystanders *E.coli* and *S. aureus* to antibiotics. While these calculations require a number of assumptions, they underscore the potentially substantial impact of vaccines on bystander selection and the need for improved data on the impact of vaccination on use of specific antimicrobials in specific populations.

## Discussion

For most bacterial species, the majority of their antibiotic exposures were the result of treatment for a condition that they did not cause. This held true across a range of different organisms and antibiotics. Carriage prevalence was the key predictive factor of the differences in proportion of bystander exposures between organisms, with species that were commonly carried asymptomatically (SI Appendix, Table S2), such as *E. coli, H. influenzae* and *S. pneumoniae*, having consistently high bystander proportions, and more rarely carried species such as *S. pyogenes* and *N. gonorrhoeae*, which are frequently associated with antibiotic-treated disease, having lower ones. Among drugs/drug classes, nitrofurantoin, used almost exclusively for urinary tract infections, had low bystander proportions for common urinary tract pathogens, which frequently are the cause of nitrofurantoin treatment. In contrast, broad-spectrum drug classes such as beta-lactams, cephalosporins, and quinolones typically have high bystander proportions for most or all species considered, because they are used for a wide variety of conditions caused by a wide variety of species, as well as for treatment of conditions that are often nonbacterial.

Quantifying bystander selection for resistance for different antibiotic-species combinations has several potential applications. As previously discussed, mathematical transmission models of antibiotic prescribing and resistance commonly assume that bystander selection is the rule rather than the exception, and these findings confirm this has been a sensible assumption, at least for outpatient antibiotic use. For policy discussions, the high bystander proportions obtained here suggest that interventions to reduce antimicrobial use may have broad effects in reducing the strength of selection across a number of bacterial species, not only the ones involved in the pathogenesis of the disease targeted by such efforts.

For example, improved adherence to guidelines on unnecessary antimicrobial prescribing might mainly affect prescribing for respiratory infections, yet might reduce selection for resistance on potential pathogens that reside on the skin (e.g. *S. aureus*) or in the gut (e.g. *E. coli* and *Klebsiella* species), as well as on respiratory bacteria. In the area of antimicrobial stewardship, these findings suggest that each reduction in inappropriate antibiotic prescribing for a particular indication may have broad impacts across many species but may not dramatically reduce the exposures to antibiotics of any one species, as long as prescribing for other indications remains unchanged.

As discussed, another example of an intervention that can reduce antimicrobial prescribing is vaccination. Vaccines can reduce the incidence of resistant infections directly (by preventing disease from their target pathogens) and indirectly (by preventing the need for antibiotic prescribing, thereby protecting bystander bacteria from exposure to antibiotics that can promote resistance). High bystander proportions are seen here for broad-spectrum antibiotic classes that are frequently prescribed for respiratory infections, and respiratory infections (including otitis media) account for a large fraction of total antimicrobial use. These considerations suggest that vaccines against pathogens that cause respiratory infections, such as *Bordetella pertussis, Streptococcus pneumoniae*, influenza virus, and respiratory syncytial virus, may substantially reduce the exposure of a broad range of pathogenic bacterial species to antibiotics, via prevention of bystander selection. Notably, this includes vaccines that prevent viral respiratory infections, which are often inappropriately treated with antibiotics (17) and perhaps prevent bacterial secondary infections that might be appropriately treated if they occurred (18). We have described an approach for using estimates of bystander exposures to estimate how vaccines could reduce exposure across various non-target pathogens. However, as noted by [REF Sevilla et al.’s perspective in this issue], quantifying the impact of vaccines on antimicrobial resistance is a complex task, and many components of such calculations will depend on the population, vaccine, and timescale considered, among other variables.

It is informative to consider the antimicrobial agents not included in our analysis. Most antimycobacterial agents have little effect on other bacterial species, while most broad-spectrum antibacterial classes are of little use against *Mycobacterium tuberculosis*. Quinolones are an exception to both rules ‐ bystander selection has been documented both in treatment of what was thought to be bacterial pneumonia but was actually tuberculosis (19) and in treatment with quinolones in a tuberculosis ward promoting the spread of quinolone-resistant *S. pneumoniae (20)*. Other exceptions include rifamycins, also commonly used for treatment of tuberculosis, and macrolides, which may be prescribed for *M. avium* complex disease. With these exceptions, bystander selection by antimycobacterial drugs is expected to be limited, and bystander selection on *M. tuberculosis* is also expected to be limited. This is reflected in an appropriate focus for tuberculosis resistance management in ensuring adequate treatment to prevent emergence of resistance and prevent transmission, rather than on bystander-focused interventions. Additionally, antiviral agents, such as the neuraminidase inhibitor oseltamivir for influenza, have no substantial known activity against other components of the (bacterial) microbiome, so the rationale for prudent use of oseltamivir would include avoiding side effects and costs, but not avoiding selection for resistance. Beyond anti-infectives, a recent *in vitro* study found that 24% of 835 therapeutic compounds with molecular targets in human cells inhibited the growth of at least one bacterial species commonly found in the human gut microbiome (21). This work suggests that our focus on antimicrobials underestimates bystander selection for resistance, but further research is needed to elucidate which drug-species combinations may be prone to such effects.

This analysis has several limitations. Firstly, all necessary inputs ‐ incidence and etiology of bacterial infections, antibiotic prescribing practices, and composition of the microbial flora ‐ are derived from different data sources and are highly heterogeneous, varying over time and by age, gender, and geographic location. Antibiotic prescribing additionally depends upon safety and toxicity profiles in certain populations. Microbiome diversity varies between and within individuals, depending on demographic characteristics, diet, and disease. For simplicity, we only consider age group stratification to calculate population-level, average estimates. Carriage prevalences and etiologies are also applied uniformly across visits. This may bias our estimates depending on the extent of microbial ecological or etiological relationships. For example, if abundance (or lack) of organism A in the microbiome contributes to the pathogenicity of organism B, organism A may be more (or less) prone to bystander exposure of antibiotics used to treat the condition caused by organism B than we calculate. Additionally, while we estimate the impact on the organism at the species level due to data constraints, selection pressures may be more relevant at the strain level; for example, true bystander selection may be lower for infrequently carried, more pathogenic strains compared to our overall, species-level estimate.

Secondly, the limitations of the datasets used in our analysis also extend to our results. For example, the HMP was conducted in a restricted, healthy study population and prevalence estimates may not be generalizable to the US population. To our knowledge, a more extensive and nationally representative source of microbiome data is unavailable, and little is known about how microbiome composition among individuals with common outpatient conditions may differ from that of healthy individuals. Additionally, HMP samples a limited number of body sites and may exclude the most relevant colonization sites for some species of interest. For example, measurement of *E. coli* in the stool as a proxy for the large intestine likely contributes to its low carriage prevalence in HMP (66.3%), leading to underestimation of colonization and thus bystander proportion. Similarly, unavailability of samples from the anterior nares for some individuals may have led to underestimation of the carriage prevalence of *S. aureus* in HMP data (12.4%). Etiologic studies are burdensome and thus often conducted among very small populations; small sample sizes may miss infrequent causative agents of disease, but this is unlikely to have a substantial effect on our point estimates. GISP, used for the analysis of *N. gonorrhoeae*, is based on convenience sampling of male patients at sexually-transmitted disease clinics. In 2016, the CDC estimated that 90.8% of patients in STD clinics received the recommended azithromycin/ceftriaxone regimen compared to 79.8% of patients in other provider settings; the same relationship, though much weaker, was observed when comparing men to women (22). Since these differences are fairly small, any underestimation of bystander selection for *N. gonorrhoeae* due to the recommended antibiotics of azithromycin and ceftriaxone is likely to be minor. Additionally, though NAMCS/NHAMCS are unique in providing a large sample of outpatient visits with corresponding diagnoses and prescriptions, a direct link between diagnosis and prescription is unavailable ‐ therefore, exposures may be incorrectly counted as “bystander”, when the prescription was in fact written for a second diagnosis caused by the species of interest. In this analysis, we make no assumptions about linkage between prescriptions and diagnoses; instead, we attribute any antibiotic prescription to all of the diagnoses recorded in the same visit. This may bias the proportion of bystander exposures in either direction, depending on the antibiotic and species pair in question; use of an alternative tiered method for assigning diagnoses to visits (37) resulted in similar findings (98% of antibiotic class and species pairs were within 5% of the values reported here).

Finally, we use prescriptions as a proxy for exposure, which is itself a proxy for selective pressure. NAMCS/NHAMCS do not contain information on whether or not the prescriptions were filled; even after being filled, we have no information on compliance to the listed medications. Furthermore, little is known about how exposures of a particular antibiotic correspond to selection pressures. This may differ widely by antibiotic, regimen, organism, and body site. Strength of selective effects at different body sites will further vary based on pharmacodynamics, pharmacokinetics, and context (e.g. microbiome composition) (25). Most of the antibiotic classes considered here may be assumed to exert some selective pressure for resistance throughout the body. Direct evidence of selection on the gut normal flora by oral antibiotics has been reported (26). Likewise, selection on the flora of the upper respiratory tract is likely the rule for many of these classes, including macrolides, penicillins, cephalosporins, and trimethoprim/sulfamethoxazole because they are routinely prescribed for upper respiratory infections and are documented to affect bacterial carriage at in the nasopharynx (27–29). Antibiotics in major classes including penicillins and cephalosporins (30), macrolides (31) and quinolones (32) have been detected in sweat, indicating that they can exert selection on skin flora. One exception to this general rule is nitrofurantoin, which tends to concentrate in the urinary tract and is accordingly used to treat UTIs. The relationship between drug concentration and selective pressure is also not straightforward, with subinhibitory concentrations likely playing an important role in selection for resistance (33). In general, the metric used in the present study, counting prescriptions as exposures, could be refined in studies of individual drugs, classes, microbes or body sites to incorporate more biological detail, and deviations from the estimates shown here will be specific to the antibiotic and organism of interest.

The bystander proportions quantified in this analysis are a step toward better characterizing the dynamics of antibiotic resistance and should be considered in the development and prioritization of interventions. In this paper, we specifically address selective pressures for resistance among potential pathogens in the microbial flora due to outpatient antibiotic prescribing. Outpatient prescribing constitutes approximately 90% of total antimicrobial volume for human health in developed countries (22, 23), but certainly further work is needed to consider the inpatient context as it affects nosocomial pathogens. In addition, off-target antibiotic exposures also contribute to resistance dynamics in other ways not captured in this analysis, including selection for resistance elements among non-pathogenic bacteria that may be horizontally transferred to pathogens and depletion of beneficial bacteria which play active roles in metabolism, pathogen resistance, and immune responses (34, 35). Research on the broader effects of antibiotic use on the microbiome is greatly needed to further understand the implications of bystander exposures on the spread of antibiotic resistance.

## Materials and methods

### Data sources

All estimates except for *Neisseria gonorrhoeae* were based on three main data sources - the National Ambulatory Medical Care Survey/National Hospital Ambulatory Medical Care Survey (NAMCS/NHAMCS) collected by the National Center for Health Statistics, the Human Microbiome Project and other studies of carriage prevalence, and etiological studies.

The NAMCS and NHAMCS are cross-sectional national surveys designed to collect data on ambulatory care services provided at office-based physician practices, emergency, and hospital outpatient departments throughout the United States. At each sampled visit, patient characteristics (e.g. age), visit characteristics (e.g. reason for visit, diagnosis, prescriptions), and physician characteristics are recorded, including up to 3 diagnoses and up to 8 prescribed medications. Sampling is based on a multistage probability sampling scheme. The most recent 2 years of fully available data for both NAMCS and NHAMCS (2010-2011) were used for this analysis. As the focus of our analysis was outpatient antibiotic use, visits that resulted in hospital or observation unit admission were excluded. Antibiotics were grouped into classes based on the Multum Lexicon system used in NAMCS/NHAMCS.

The first phase of the Human Microbiome Project (HMP) consisted of collecting microbiome samples from 300 healthy individuals between the ages of 18 and 40 at multiple timepoints across five major body sites: the nasal passages, oral cavity, skin, gastrointestinal tract, and urogenital tract. Microbial composition was characterized using MetaPhlAn2 (36), a taxonomic profiling method for whole-metagenomic shotgun samples. Prevalence estimates from HMP data were based on presence of the species at any body site. For children under 5 years old, carriage prevalences were compiled from primary sources in the literature (SI Appendix, Table S1). As individual carriage studies tended to collect samples from only one body site, carriage prevalences at each body site were estimated as an average across studies weighted by sample size, and overall prevalence was calculated assuming independence at each body site. This process was also used to estimate carriage prevalences of *S. pyogenes* and *S. pneumoniae* in the >5 age group, as MetaPhlAn2 did not distinguish between these and closely related species (e.g. *S. mitis, S. oralis)*. Etiologies for conditions of interest were based on etiologic studies cited in the medical resource UpToDate (SI Appendix, Table S3).

### Calculations of bystander proportions

A bystander exposure was defined as a prescription of antibiotic (or antibiotic class) *a* received by an individual carrying species s for a diagnosis of condition *c* that was not caused by *s*. Exposures were estimated on average at the population level. Let *B*_*as*_ be the proportion of bystander exposures of antibiotic *a* received by species *s*, equivalent to one minus the ratio of *N*_*as*_, the number of exposures of antibiotic *a* received by species s for a case of some condition *c* that was caused by species *s*, and *T*_*as*_, the total number of exposures of antibiotic *a* received by species *s*. Additionally, let *d*_*acg*_ be the number of prescriptions of antibiotic *a* written for condition *c* in age group *g*, let *p*_*scg*_ be the proportion of cases of condition *c* who are colonized with species *s* in age group *g*, and let *e*_*scg*_ be the proportion of cases of condition *c* caused by species s in age group g. The proportion *p*_*scg*_ was calculated under the assumption that all individuals with condition *c* not caused by species *s* were colonized with species s at the group-specific prevalence estimated from HMP or other studies (denoted *p*_*sg*_), while individuals with condition c caused by species s were colonized with species s with probability 1: *p*_*scg*_ = *e*_*scg*_ + (1-*e*_*scg*_)*p*_*sg*_. Since the inputs *d*_*acg*_, *e*_*scg*_, and *p*_*sg*_ may be highly variable by age, estimates were summed over three age strata *g* (<1 year old, 1-5 years old, and over 5 years old). The proportion of bystander exposures for antibiotic *a*, species *s*, and condition *c* was calculated as follows:

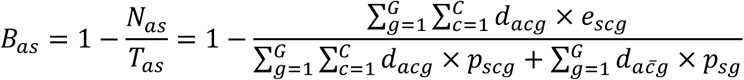

Conditions were based on diagnostic categories delineated by Fleming-Dutra et al. (37) with the following exceptions: 1) “Other bacterial infections” includes all codes listed under “miscellaneous bacterial infections” plus other intestinal infectious diseases (ICD-9CM codes: 001-008), but excludes a subset of infectious diseases (040-041, 130-139), mastoiditis (383), and peritonsillar abscess (475); 2) we include only cellulitis (681-682) from the category “Skin, cutaneous and mucosal infections”. The set of conditions *C* includes conditions for which antibiotic use was relatively high (>2% of weighted prescriptions; viral upper respiratory tract infection contributed the most, at 10.8% of weighted prescriptions) and reasonable estimates of *e*_*scg*_ were available. When diagnoses were excluded, this was most often due to one of these two limitations. Influenza was also included due to clear etiology and vaccination-related interest, and acne was included due to high tetracycline use for this indication. All cases with ICD-9CM code indicating the causative agent (e.g. 480: viral pneumonia, 481: streptococcal pneumonia [*Streptococcus pneumoniae* pneumonia]) were attributed to that agent.

The second term in the denominator of the proportion of bystander exposures accounts for exposures of antibiotic *a* that were not associated with any of the conditions in *C*, where *d*_*ac̄g*_ represents prescriptions of antibiotic *a* that occur at visits not associated with any condition in C. The use of *p*_*sg*_ in this term implies that our species of interest are rarely, if ever, causative agents for conditions that are not included in our analysis. Violations of this assumption will lead to overestimation of the proportion of bystander exposures. For comparison, we performed the analysis excluding this term (SI Appendix, Figure S5) as lower bound estimates of the proportion of bystander exposures for all species-antibiotic class pairs.

Confidence intervals were estimated by simulation. The proportion of bystander exposures was calculated for 1000 random draws of *d*_*acg*_, *p*_*sg*_, and *e*_*scg*_ based on empirically estimated distributions. Draws for *d*_*acg*_ were based on the normal distribution, using variances calculated by the ‘survey’ package in R (38). For HMP prevalence estimates, with A presences and B absences, random draws were simulated from a beta distribution with parameters (*A* + 0.5, *B* + 0.5), the posterior distribution using Jeffreys prior.

Resampling was done similarly for etiological fractions. The 2.5^th^ and 97.5^th^ percentiles were utilized as the bounds of the 95% confidence interval. Table 1 provides a summary of notation used in this analysis.

**Table 1.** Summary of notation with definitions, sources, and relevant figures and tables.

| Variable | Definition | Source | Figure/Table |
| --- | --- | --- | --- |
| $d_{acg}$ | Number of prescriptions (using nationally representative weights) of antibiotic $a$ associated with condition $c$ in age group $g$ | NAMCS/NHAMCS 2010-11 | Figure S2 (unweighted) |
| $d_{a\bar{c}g}$ | Number of prescriptions (using nationally representative weights) of antibiotic $a$ associated with all conditions not explicitly included in analysis ($\bar{c}$) in age group $g$ | NAMCS/NHAMCS 2010-11 | N/A |
| $p_{sg}$ | Carriage prevalence of species $s$ in age group $g$ | HMP and carriage studies | Figure 1, Table S2 |
| $p_{scg}$ | Carriage prevalence of species $s$ among individuals with condition $c$ in age group $g$ | $e_{scg} + (1 - e_{scg})p_{sg}$ | N/A |
| $e_{scg}$ | Proportion of cases of condition $c$ in age group $g$ caused by species $s$ | Etiologic studies | Figure 1, Table S3 |
| $N_{as}$ | Number of exposures of antibiotic $a$ experienced by species $s$ for cases of conditions in $C$ caused by species $s$ | $\sum_{g=1}^G \sum_{c=1}^C d_{acg} \times e_{scg}$ | N/A |
| $T_{as}$ | Total number of exposures of antibiotic $a$ experienced by species $s$ over all conditions | $\sum_{g=1}^G \sum_{c=1}^C d_{acg} \times p_{scg} + \sum_{g=1}^G d_{a\bar{c}g} \times p_{sg}$ | N/A |
| $B_{as}$ | Proportion of bystander exposures for antibiotic $a$ and species $s$ | $1 - \frac{N_{as}}{T_{as}}$ | Figures 1, S1 and S4 |

*Calculations of bystander proportion for* N. gonorrhoeae. Separate analyses were conducted for *N. gonorrhoeae* using additional data from the Gonococcal Isolate Surveillance Project (GISP) (14). The same formula for the proportion of bystander exposures was applied. When GISP data was reported by clinic site, the weighted average by sample size over all clinics was used. Age groups were further stratified into those used by GISP (<20, 20-24, 25-29, 30-34, 35-39, 40-44, >45 years old) for this analysis.

*N. gonorrhoeae* was assumed to be the causative agent for all gonorrhea cases only (*e*_*scg*_=1 for gonorrhea and 0 for all other conditions of interest). The number of prescriptions of antibiotic *a* written for all conditions except gonorrhea was calculated from NAMCS/NHAMCS as in the previous analyses. We include only prescriptions for patients >5 years old, as we expect gonococcal infection and carriage to be very rare among children age 5 and below. For gonorrhea, the number of prescriptions of antibiotic *a* written in 2010-2011 was estimated by multiplying the total number of reported gonorrhea cases (309,341 in 2010 (39) and 321,849 in 2011 (40)) by the proportion of GISP participants treated with *a*.

Prescriptions were counted equally regardless of dosage. Data on quinolone and cephalosporin prescribing only were available for 2010, while 2011 reports also included macrolides and tetracyclines. Therefore, bystander proportions for azithromycin and doxycycline are only available for 2011.

*N. gonorrhoeae* is identified in HMP, but the measured prevalence of 37%, with 95% of these identified in oral isolates, indicates that this data may include false positives. Miller et al. report that prevalence of *N. gonorrhoeae* in urine samples from a nationally representative sample of young adults (National Longitudinal Study of Adolescent Health) was 0.64% among participants 20-21 years old, 0.47% among participants 22-23 years old, and 0.24% among participants 24-25 years old. Using the target population weights reported in Table 1 of the same paper, we estimated the carriage prevalence of *N. gonorrhoeae* to be 0.56% among participants 20-23 years old and 0.48% among participants 20-25 years old (41). We applied the mean of these two groups, 0.52%, as the carriage prevalence of *N. gonorrhoeae* in the 20-24 age category used by GISP. For all other age groups designated by GISP, this prevalence was inflated by the relative proportion of GISP isolates from that age group. For example, in 2010, 31.4% of GISP isolates were sampled from individuals aged 20-24 years old, while 21.2% were from individuals aged 25-29 years old. The carriage prevalence among 20-24 year olds was multiplied by 21.2%/31.4% to estimate carriage prevalence in the 25-29 age category.

### Impact of vaccine

To approximate the impact of a vaccine in reducing antimicrobial exposure of nontargeted species (e.g. *E. coli* for a pneumococcal vaccine), we initially assume as an input the observed reduction *r* in all-cause antimicrobial use in a particular age group, such as the 5.4% reduction in all-cause antibiotic use in a randomized pneumococcal conjugate vaccine trial in 0-2 year-olds (5). From our analysis, we approximate values for 0-2 year-olds as the average of results from the 0-1 and 1-5 age groups. We reason as follows:

Table 2 shows the possible combinations of presence/absence of E. coli in a treated patient, and E. coli as cause or not cause of the treatment. One cell (absent, but causal) is empty because by assumption the species must be present to cause treatment. Let *A*, *B*, and *D* represent proportions of all treatments so *A* + *B* + *D* = 1.

**Table 2.** Classifying all-cause antibiotic treatments with respect to a potential bystander species, *E. coli*, as present or not, and cause of treatment or not.

|  | *E. coli* present in treated patient |  |  |
| --- | --- | --- | --- |
| *E. coli* is the cause of treatment |  | - | + |
| | - | $A$ | $B$ |
| | + | | $D$ |

In our example, the total treatment reduction, *r*, is 5.4% of all treatments. However, this reduction is unequally apportioned. All of the reduction occurs in categories *A* and *B*, because we assume that PCV would have no effect on the rate of treatment for a disease that was caused by *E. coli*. Define *p*_*Ec*_ as the prevalence of E. coli in the microbiome data for the relevant age group. Then by our modeling assumptions, 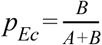. Thus, the amount of treatment reduction in category B is 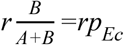.

We seek the proportional reduction in *B* + *D*, the exposure of *E. coli* to treatment. D is unchanged, so the reduction is 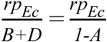. Defining the proportion of bystander exposures for *E. coli* to all antibiotics as 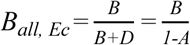 some algebra yields the quantity we seek, the reduction in *E. coli’s* total (causal plus bystander) exposure to antibiotics attributable to a reduction *r* in all-cause antibiotic treatment from a vaccine that prevents no disease caused by *E. coli*:

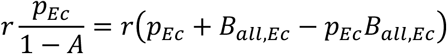

Analogous calculations can be made for any other bacterial species for which disease is not reduced by the vaccine. For pathogens (e.g. *H. influenzae)* in which vaccination may cause a reduction in the amount of disease they cause (e.g. through indirectly preventing non-pneumococcal otitis media (15, 16)), this estimate would be a lower bound.

All data for this project is publicly available. Code will be made available on Github.

## Acknowledgements

We thank Curtis Huttenhower, Eric Franzosa, and Jason Lloyd-Price for helpful comments and assistance with MetaPhlAn2 data. This work was supported by Grant U54GM088558 (Models of Infectious Disease Agent Study [MIDAS] - Center for Communicable Disease Dynamics) from the National Institute of General Medical Sciences, Grant R01 AI132606 from the National Institute of Allergy and Infectious Diseases (Y.H.G.), Grant CK000538-01 from the Centers for Disease Control and Prevention, and the Doris Duke Charitable Foundation. The content is solely the responsibility of the authors and does not necessarily represent the official views of the National Institute of General Medical Sciences, National Institutes of Health, National Institute of Allergy and Infectious Diseases, Centers for Disease Control and Prevention, Department of Health and Human Services, or the Doris Duke Charitable Foundation.

## Supplementary Information (SI)

**Figure S1.**
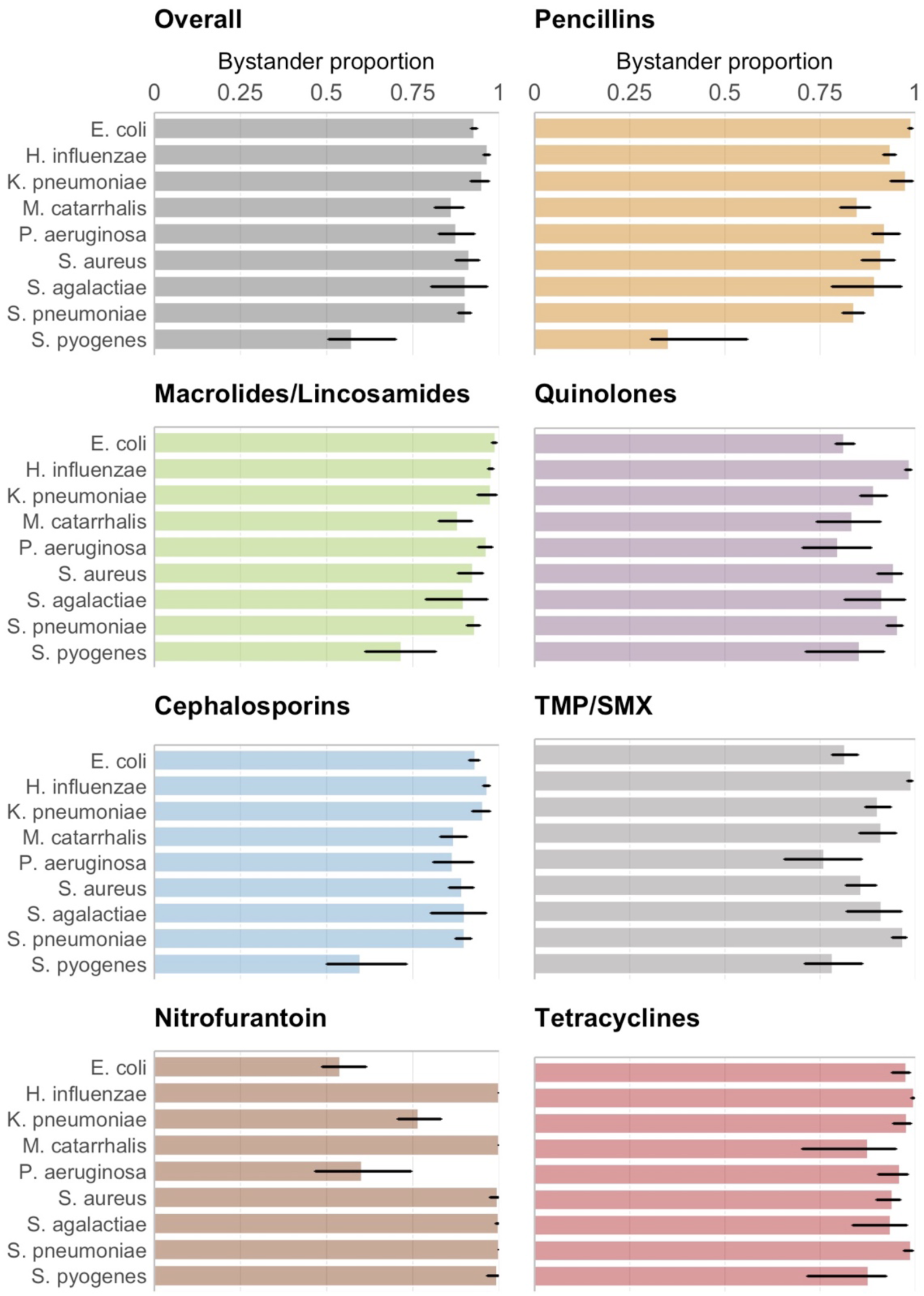
Proportion of bystander exposures (with 95% confidence interval) by antibiotic class and species. “Overall” estimates reflect exposures to antibiotics in any of the included classes. Results for TMP/SMX and nitrofurantoin are for the individual drug instead of an antibiotic class.

**Figure S2.**
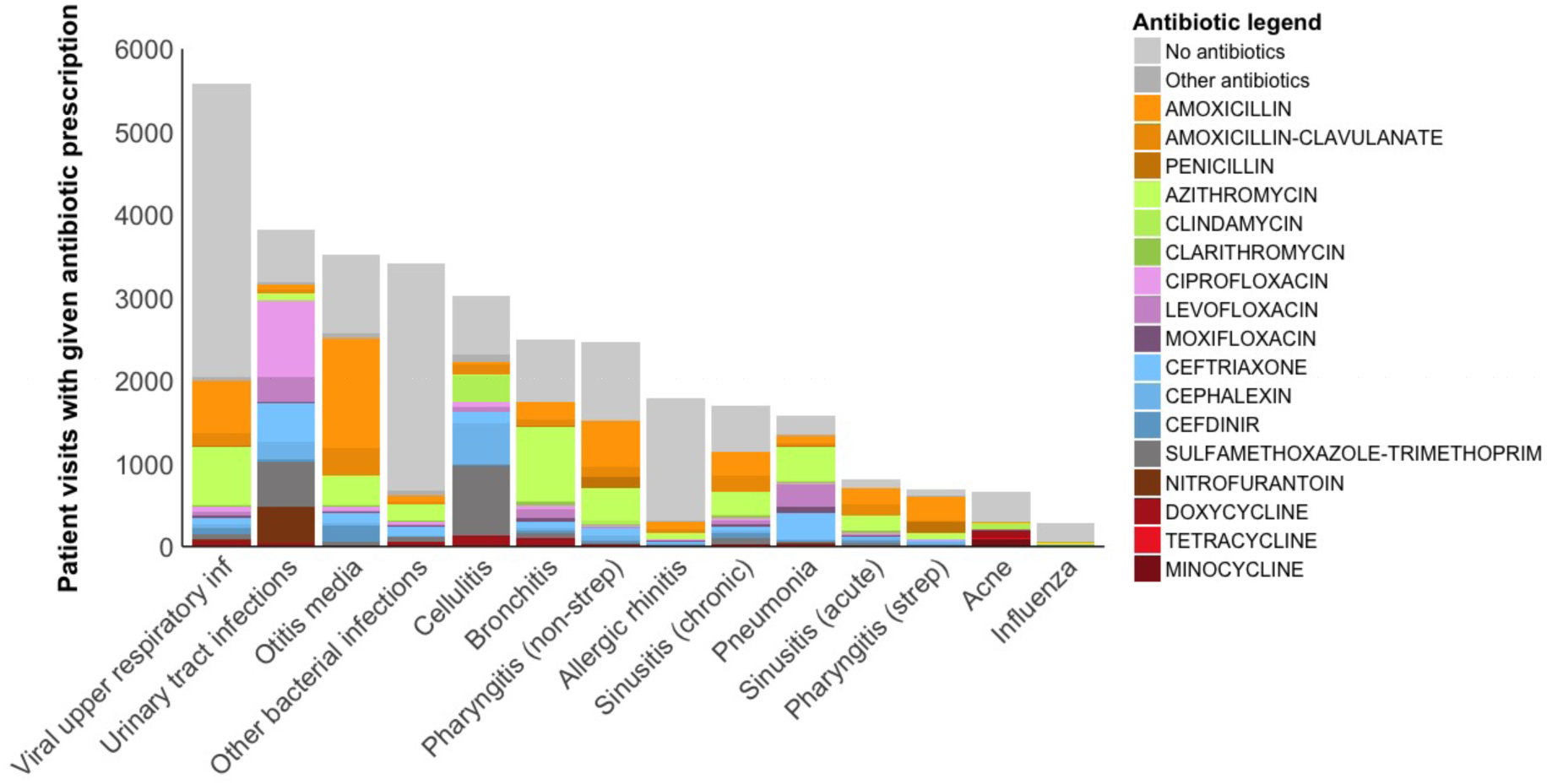
Number of sampled outpatient visits (unweighted) from NAMCS/NHAMCS 2010-2011 with given diagnosis and antibiotic prescription.

**Figure S3.**
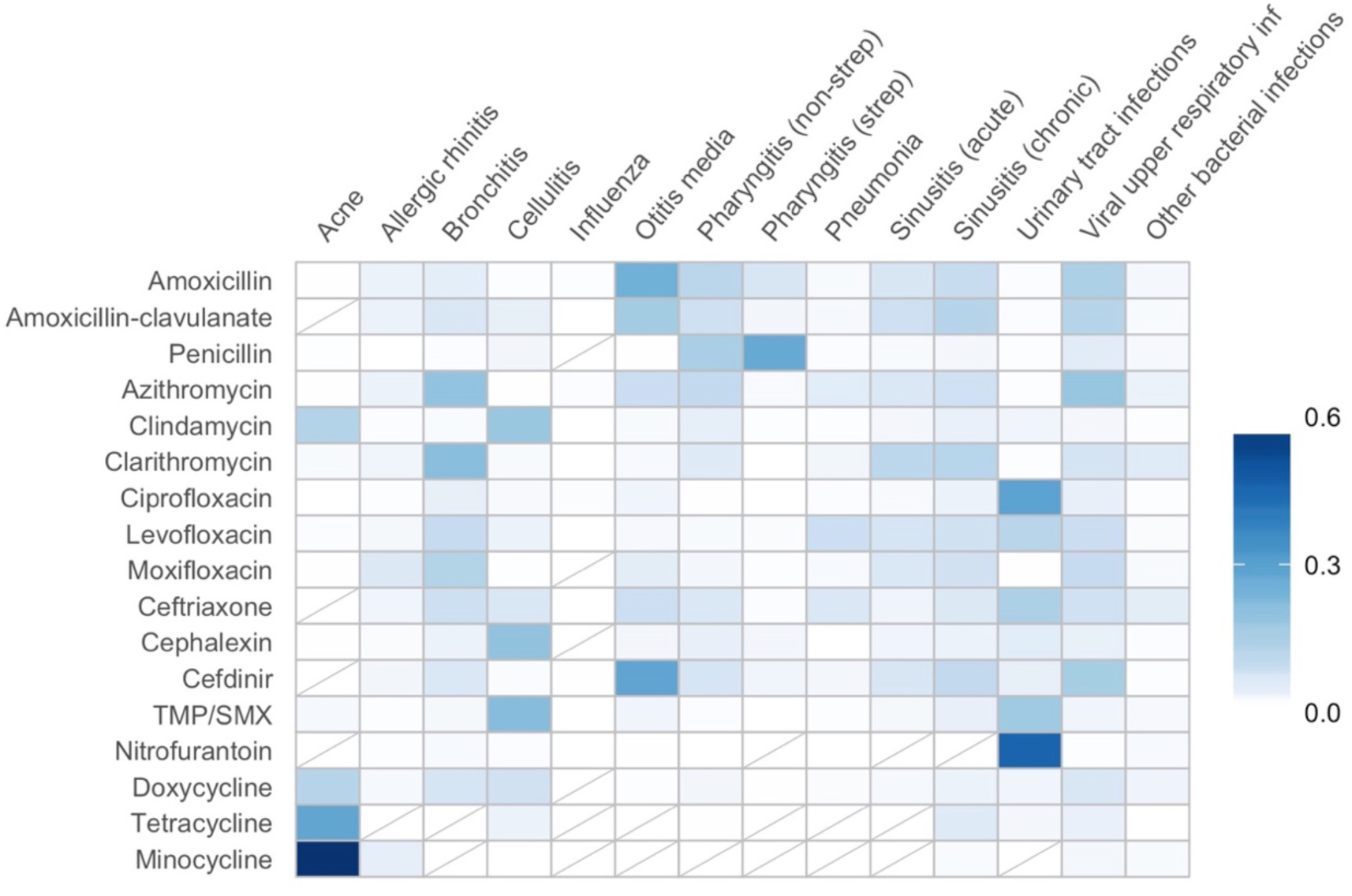
Heat map shading represents the proportion of visits (after weighting to be nationally representative) with a diagnosis of the specified condition, given that the visit resulted in a prescription of the specified antibiotic. Rows are not required to sum to 100% as only a subset of conditions are shown, and each visit may be associated with more than one condition.

**Figure S4.**
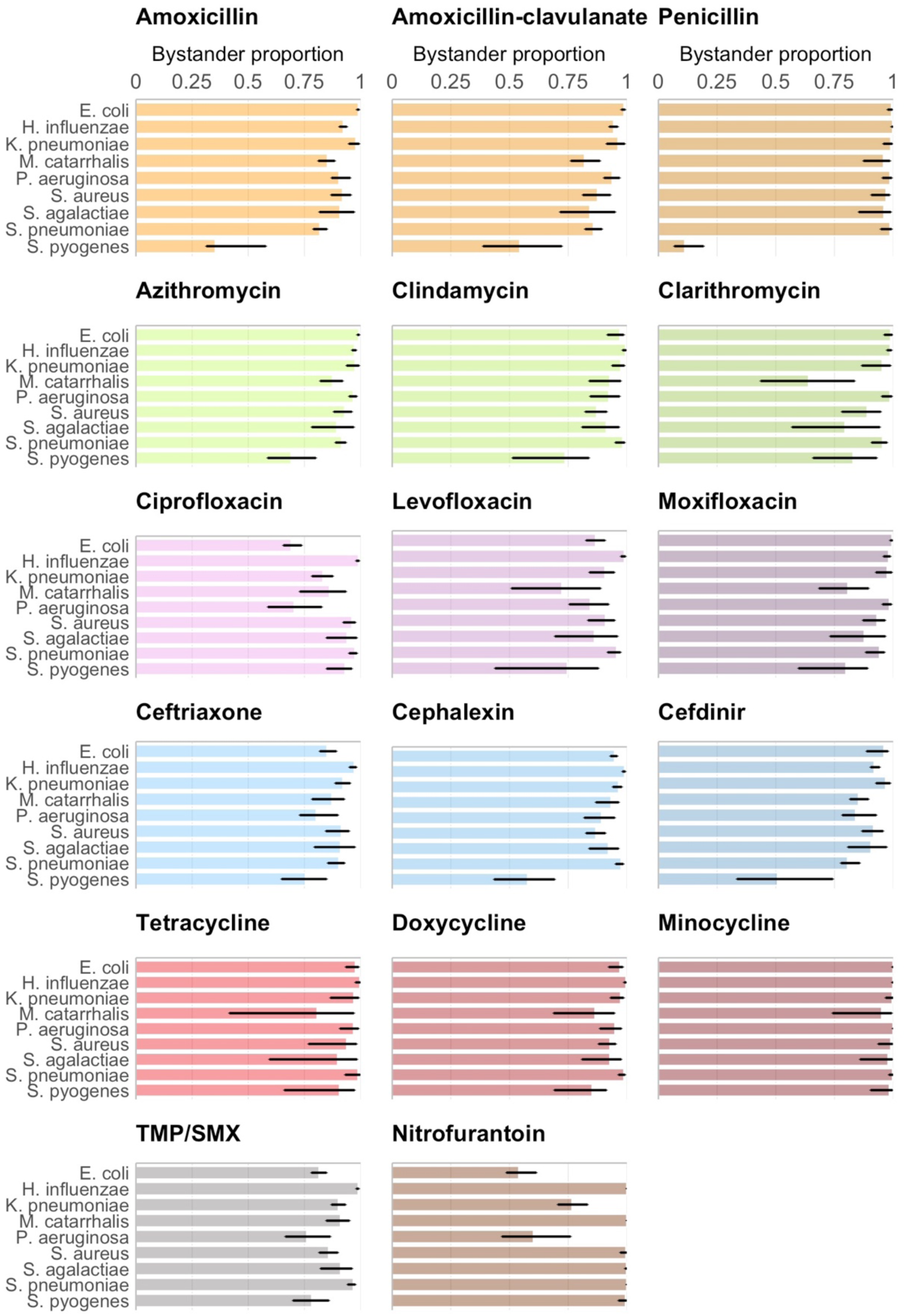
Proportion of bystander exposures (with 95% confidence interval) by antibiotic and species.

**Figure S5.**
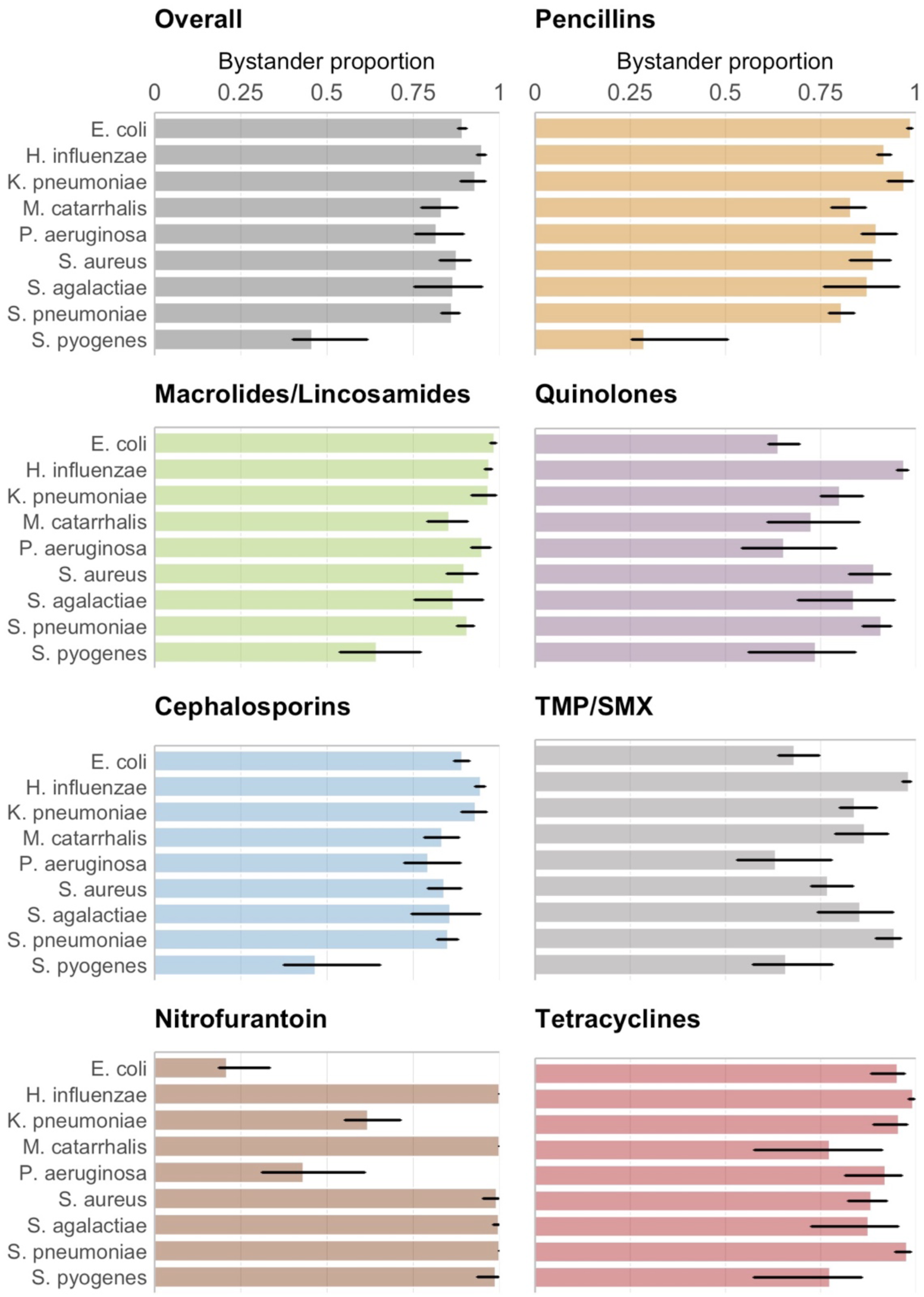
Proportion of bystander exposures (with 95% confidence interval) by antibiotic class and species, excluding the term 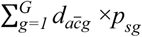 from the denominator. “Overall” estimates reflect exposures to antibiotics in any of the included classes. Results for TMP/SMX and nitrofurantoin are for the individual drug instead of an antibiotic class.

**Table S1.** Carriage studies used to characterize microbial prevalences for which HMP data was unavailable. In addition to prevalences among children <5 years old, additional carriage studies were also used for *S. pyogenes* and *S. pneumoniae* in the >5-year-old age group as taxonomic profiling of HMP data via MetaPhlAn2 does not distinguish between these and similar species. Specific studies were not identified for *P. aeruginosa* and *S. agalactiae* for children from 1 to 5 years old; the prevalences among children under 1 year old were imputed in these cases.

| Article | Age group | Body site | Organisms |
| --- | --- | --- | --- |
| Bäckhed et al. 2015 (1) | <1 year old | Gastrointestinal | *P. aeruginosa*<br>*S. agalactiae* |
| Bogaert et al. 2011 (2) | 1-5 years old | Nasopharyngeal | *H. influenzae* |
| Mainous et al. 2006 (3) | 1-5 years old | Nasopharyngeal | *S. aureus* |
| Regev-Yochay et al. 2004 (4) | <1 year old<br>1-5 years old | Nasopharyngeal | *S. aureus*<br>*S. pneumoniae* |
| Verhaegh et al. 2010 (5) | <1 year old<br>1-5 years old | Nasopharyngeal | *M. catarrhalis* |
| Pettigrew et al. 2012 (6) | <1 year old<br>1-5 years old | Upper respiratory tract | *H. influenzae*<br>*M. catarrhalis*<br>*S. pneumoniae* |
| Holgersen et al. 2015 (7) | <1 year old<br>1-5 years old | Oral | *E. coli*<br>*H. influenzae*<br>*K. pneumoniae*<br>*S. aureus*<br>*S. pyogenes* |
| Yassour et al. 2016 (8)<br>(DIABIMMUNE cohort) | <1 year old<br>1-5 years old | Gastrointestinal | *E. coli*<br>*H. influenzae*<br>*K. pneumoniae*<br>*S. aureus* |
| Ginsburg et al. 1985 (9) | All | Throat | *S. pyogenes* |
| Gunnarsson et al. 1997 (10) | All | Throat | *S. pyogenes* |
| Hammitt et al. 2006 (11) | All | Nasopharyngeal | *S. pneumoniae* |
| Huang et al. 2009 (12) | All | Nasopharyngeal | *S. pneumoniae* |

**Table S2.**
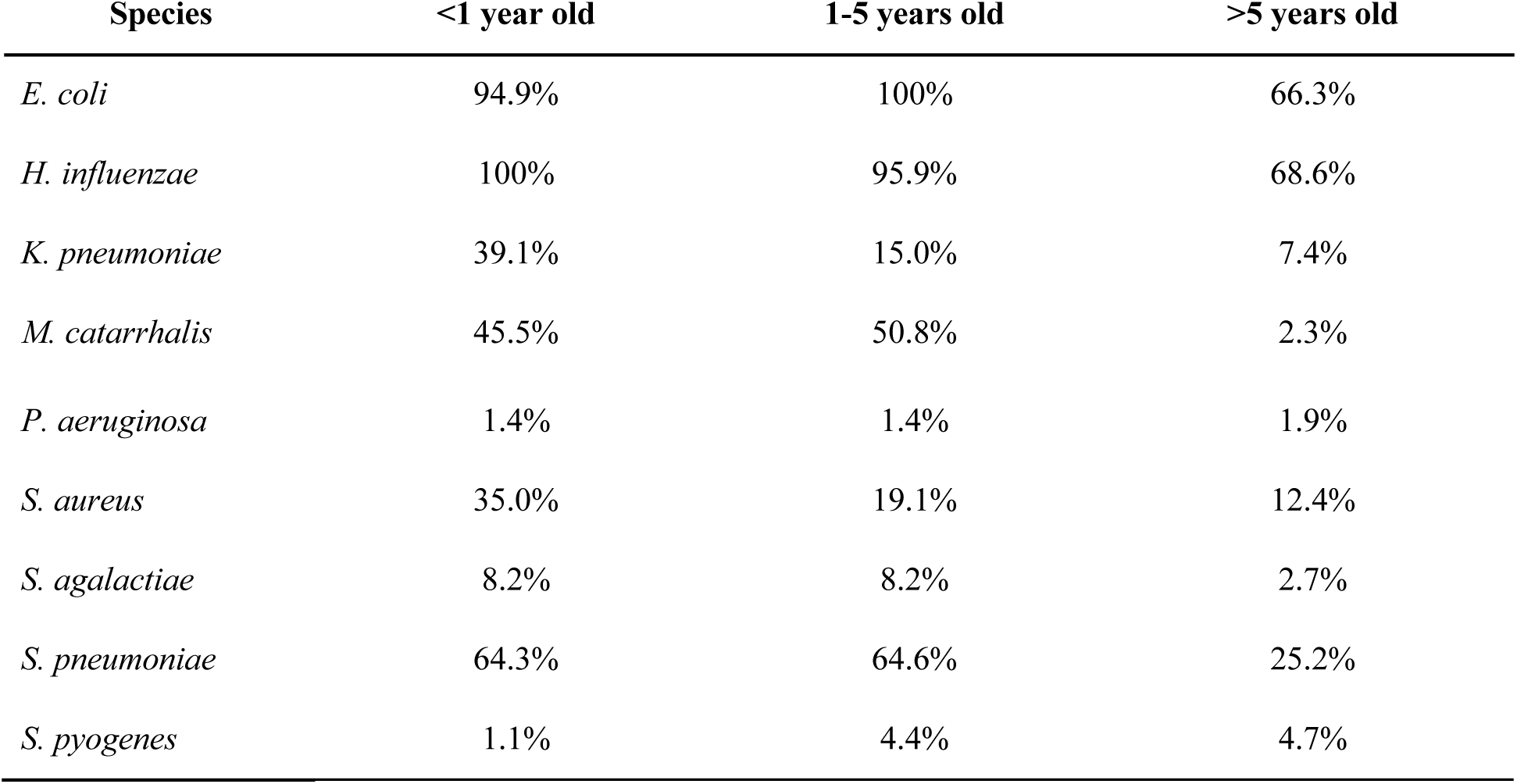
Carriage prevalence estimates by age group and species from the Human Microbiome Project (HMP) and carriage studies.

| Species | <1 year old | 1-5 years old | >5 years old |
| --- | --- | --- | --- |
| *E. coli* | 94.9% | 100% | 66.3% |
| *H. influenzae* | 100% | 95.9% | 68.6% |
| *K. pneumoniae* | 39.1% | 15.0% | 7.4% |
| *M. catarrhalis* | 45.5% | 50.8% | 2.3% |
| *P. aeruginosa* | 1.4% | 1.4% | 1.9% |
| *S. aureus* | 35.0% | 19.1% | 12.4% |
| *S. agalactiae* | 8.2% | 8.2% | 2.7% |
| *S. pneumoniae* | 64.3% | 64.6% | 25.2% |
| *S. pyogenes* | 1.1% | 4.4% | 4.7% |

**Table S3.**
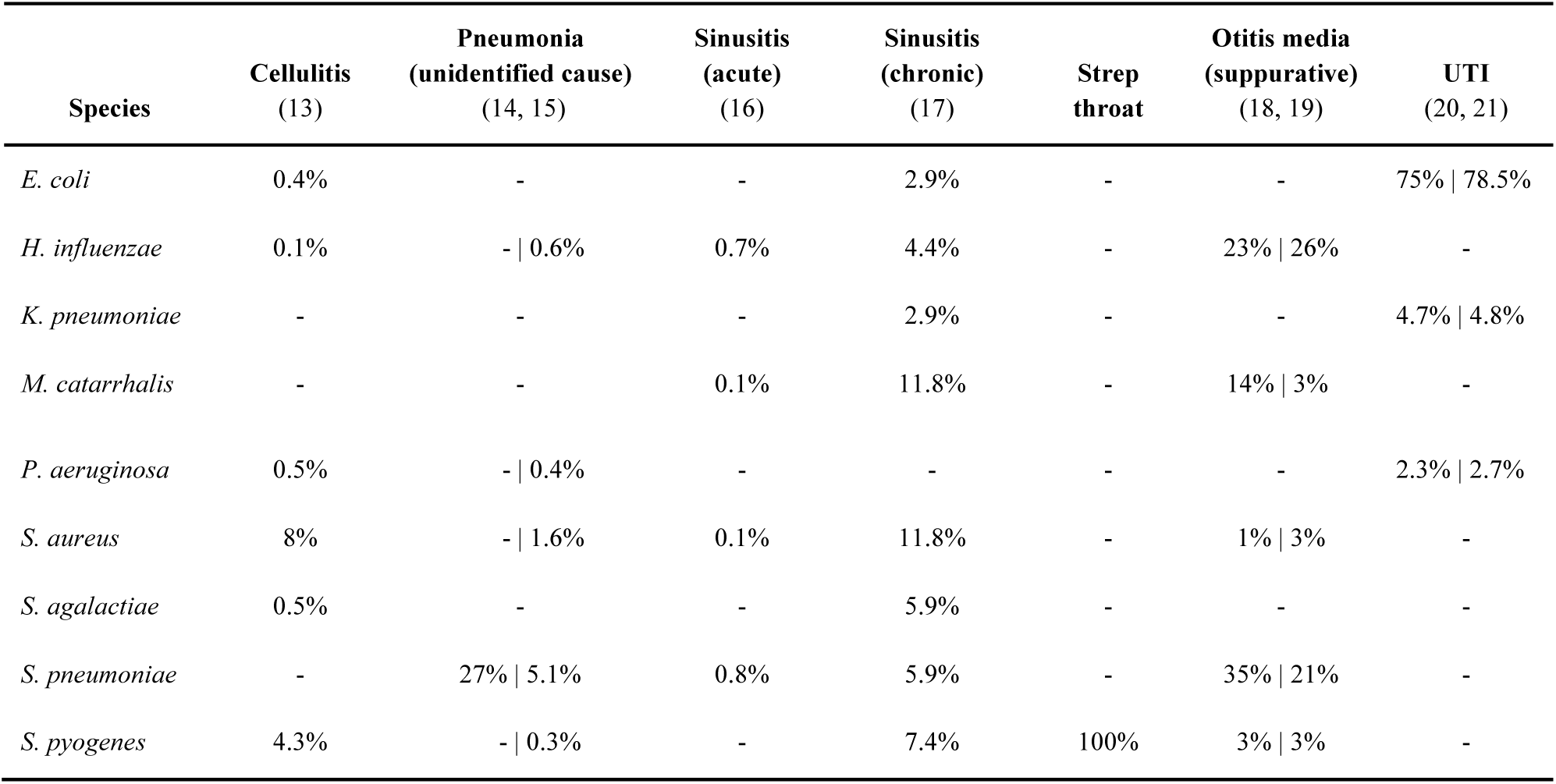
Estimated etiologies by condition. Conditions in which none of our species of interest are causative agents are excluded. If two numbers are shown, the number to the left was applied to children under 5 years old, and the number to the right was applied to individuals over 5. Diagnoses with etiology specified by ICD-9CM code (e.g. 481: pneumococcal pneumonia) were attributed to the appropriate organism.

## References

1. Sullivan A, Edlund C, Nord CE (2001) Effect of antimicrobial agents on the ecological balance of human microflora. Lancet Infect Dis 1(2): 101–114.

2. Gustafsson I, et al. (2003) Bacteria with increased mutation frequency and antibiotic resistance are enriched in the commensal flora of patients with high antibiotic usage. J Antimicrob Chemother 52(4):645–650.

3. Lindgren M, Löfmark S, Edlund C, Huovinen P, Jalava J (2009) Prolonged impact of a one-week course of clindamycin on Enterococcus spp. in human normal microbiota. Scand J Infect Dis 41(3):215–219.

4. Nyberg SD, et al. (2007) Long-term antimicrobial resistance in Escherichia coli from human intestinal microbiota after administration of clindamycin. Scand J Infect Dis 39(6-7):514–520.

5. Fireman B, et al. (2003) Impact of the pneumococcal conjugate vaccine on otitis media. Pediatr Infect Dis J 22(1): 10–16.

6. Lau WCY, et al. (2015) Impact of pneumococcal conjugate vaccines on childhood otitis media in the United Kingdom. Vaccine 33(39):5072–5079.

7. Cobey S, et al. (2017) Host population structure and treatment frequency maintain balancing selection on drug resistance. J R Soc Interface 14(133). doi: 10.1098/rsif.2017.0295.

8. Bonhoeffer S, Lipsitch M, Levin BR (1997) Evaluating treatment protocols to prevent antibiotic resistance. Proc Natl Acad Sci U S A 94(22): 12106–12111.

9. Davies NG, Flasche S, Jit M, Atkins KE (2017) Within-host dynamics explain the coexistence of antibiotic-sensitive and resistant bacteria. bioRxiv. Available at: https://www.biorxiv.org/content/early/2017/11/10/217232.abstract.

10. Atkins KE, et al. (2017) Use of mathematical modelling to assess the impact of vaccines on antibiotic resistance. Lancet Infect Dis. doi: 10.1016/S1473-3099(17)30478-4.

11. Lipsitch M, Siber GR (2016) How Can Vaccines Contribute to Solving the Antimicrobial Resistance Problem? MBio 7(3). doi:10.1128/mBio.00428-16.

12. Olesen SW, et al. (2018) Azithromycin susceptibility in Neisseria gonorrhoeae and seasonal macrolide use. J Infect Dis. doi:10.1093/infdis/jiy551.

13. Wind CM, et al. (2017) Decreased azithromycin susceptibility of Neisseria gonorrhoeae isolates in patients recently treated with azithromycin. Clin Infect Dis. doi: 10.1093/clinid/cix249.

14. CDC (2018) GISP - Gonorrhea - STD information from CDC. Available at: https://www.cdc.gov/std/gisp/default.htm [Accessed June 12, 2018].

15. Dagan R, Pelton S, Bakaletz L, Cohen R (2016) Prevention of early episodes of otitis media by pneumococcal vaccines might reduce progression to complex disease. Lancet Infect Dis 16(4):480–492.

16. Ben-Shimol S, et al. (2016) Impact of Widespread Introduction of Pneumococcal Conjugate Vaccines on Pneumococcal and Nonpneumococcal Otitis Media. Clin Infect Dis 63(5):611–618.

17. Goyal MK, et al. (2017) Racial and Ethnic Differences in Antibiotic Use for Viral Illness in Emergency Departments. Pediatrics 140(4). doi: 10.1542/peds.2017-0203.

18. Loeb M (2010) Community-acquired pneumonia. BMJ Clin Evid 2010. Available at: https://www.ncbi.nlm.nih.gov/pubmed/21418681.

19. Shen G-H, et al. (2012) Does empirical treatment of community-acquired pneumonia with fluoroquinolones delay tuberculosis treatment and result in fluoroquinolone resistance in Mycobacterium tuberculosis? Controversies and solutions. Int J Antimicrob Agents 39(3):201–205.

20. von Gottberg A, et al. (2008) Emergence of levofloxacin-non-susceptible Streptococcus pneumoniae and treatment for multidrug-resistant tuberculosis in children in South Africa: a cohort observational surveillance study. Lancet 371(9618): 1108–1113.

21. Maier L, et al. (2018) Extensive impact of non-antibiotic drugs on human gut bacteria. Nature. doi: 10.1038/nature25979.

22. Weston EJ (2018) Adherence to CDC Recommendations for the Treatment of Uncomplicated Gonorrhea — STD Surveillance Network, United States, 2016. MMWR Morb Mortal Wkly Rep 67. doi: 10.15585/mmwr.mm6716a4.

23. Swedres-Svarm (2014) Consumption of antibiotics and occurrence of antibiotic resistance in Sweden (Public Health Agency of Sweden, National Veterinary Institute).

24. Canadian Antimicrobial Resistance Surveillance System 2017 Report (2017) (Public Health Agency of Canada).

25. Ferrer M, Méndez-García C, Rojo D, Barbas C, Moya A (2017) Antibiotic use and microbiome function. Biochem Pharmacol 134:114–126.

26. Lange K, Buerger M, Stallmach A, Bruns T (2016) Effects of Antibiotics on Gut Microbiota. Dig Dis 34(3):260–268.

27. Dagan R, Barkai G, Leibovitz E, Dreifuss E, Greenberg D (2006) Will reduction of antibiotic use reduce antibiotic resistance?: The pneumococcus paradigm. Pediatr Infect Dis J 25(10):981–986.

28. Samore MH, et al. (2006) Mechanisms by which antibiotics promote dissemination of resistant pneumococci in human populations. Am J Epidemiol 163(2): 160–170.

29. Feikin DR, et al. (2000) Increased carriage of trimethoprim/sulfamethoxazole-resistant Streptococcus pneumoniae in Malawian children after treatment for malaria with sulfadoxine/pyrimethamine. J Infect Dis 181(4): 1501–1505.

30. Høiby N, Pers C, Johansen HK, Hansen H, The Copenhagen Study Group on Antibiotics in Sweat (2000) Excretion of β-Lactam Antibiotics in Sweat—a Neglected Mechanism for Development of Antibiotic Resistance? Antimicrob Agents Chemother 44(10):2855–2857.

31. Strausbaugh LJ, Bolton WK, Dilworth JA, Guerrant RL, Sande MA (1976) Comparative pharmacology of josamycin and erythromycin stearate. Antimicrob Agents Chemother 10(3):450–456.

32. Høiby N, et al. (1997) Excretion of ciprofloxacin in sweat and multiresistant Staphylococcus epidermidis. Lancet 349(9046):167–169.

33. Andersson DI, Hughes D (2014) Microbiological effects of sublethal levels of antibiotics. Nat Rev Microbiol 12(7):465–478.

34. Dethlefsen L, Relman DA (2011) Incomplete recovery and individualized responses of the human distal gut microbiota to repeated antibiotic perturbation. Proc Natl Acad Sci U S A 108 Suppl 1:4554–4561.

35. Dethlefsen L, Huse S, Sogin ML, Relman DA (2008) The pervasive effects of an antibiotic on the human gut microbiota, as revealed by deep 16S rRNA sequencing. PLoSBiol 6(11):e280.

36. Truong DT, et al. (2015) MetaPhlAn2 for enhanced metagenomic taxonomic profiling. Nat Methods 12(10):902–903.

37. Fleming-Dutra KE, et al. (2016) Prevalence of Inappropriate Antibiotic Prescriptions Among US Ambulatory Care Visits, 2010-2011. JAMA 315(17):1864–1873.

38. Lumley T (2016) survey: analysis of complex survey samples.

39. CDC - STD Surveillance, 2010 - Gonorrhea Available at: https://www.cdc.gov/std/stats10/gonorrhea.htm [Accessed June 12, 2018].

40. CDC - STD Surveillance, 2011 - Gonorrhea Available at: https://www.cdc.gov/std/stats11/gonorrhea.htm [Accessed June 12, 2018].

41. Miller WC (2004) Prevalence of Chlamydial and Gonococcal Infections Among Young Adults in the United States. JAMA 291(18):2229.

## References

1. Bäckhed F, et al. (2015) Dynamics and Stabilization of the Human Gut Microbiome during the First Year of Life. Cell Host Microbe 17(5):690–703.

2. Bogaert D, et al. (2011) Variability and diversity of nasopharyngeal microbiota in children: a metagenomic analysis. PLoS One 6(2):e17035.

3. Mainous AG 3rd, Hueston WJ, Everett CJ, Diaz VA (2006) Nasal carriage of Staphylococcus aureus and methicillin-resistant S aureus in the United States, 2001-2002. Ann Fam Med 4(2): 132–137.

4. Regev-Yochay G, et al. (2004) Association between carriage of Streptococcus pneumoniae and Staphylococcus aureus in Children. JAMA 292(6):716–720.

5. Verhaegh SJC, et al. (2010) Determinants of Moraxella catarrhalis colonization in healthy Dutch children during the first 14 months of life. Clin Microbiol Infect 16(7):992–997.

6. Pettigrew MM, et al. (2012) Upper respiratory tract microbial communities, acute otitis media pathogens, and antibiotic use in healthy and sick children. Appl Environ Microbiol 78(17):6262–6270.

7. Lif Holgerson P, Holgerson PL, Öhman C, Rönnlund A, Johansson I (2015) Maturation of Oral Microbiota in Children with or without Dental Caries. PLoS One 10(5):e0128534.

8. Yassour M, et al. (2016) Natural history of the infant gut microbiome and impact of antibiotic treatment on bacterial strain diversity and stability. Sci Transl Med 8(343):343ra81.

9. Ginsburg CM, et al. (1985) Seroepidemiology of the group-A streptococcal carriage state in a private pediatric practice. Am J Dis Child 139(6):614–617.

10. Gunnarsson RK, Holm SE, Söderström M (1997) The prevalence of beta-haemolytic streptococci in throat specimens from healthy children and adults. Implications for the clinical value of throat cultures. Scand J Prim Health Care 15(3): 149–155.

11. Hammitt LL, et al. (2006) Indirect effect of conjugate vaccine on adult carriage of Streptococcus pneumoniae: an explanation of trends in invasive pneumococcal disease. J Infect Dis 193(11): 1487–1494.

12. Huang SS, et al. (2009) Continued Impact of Pneumococcal Conjugate Vaccine on Carriage in Young Children. Pediatrics 124(1):e1–e11.

13. Chira S, Miller LG (2010) Staphylococcus aureus is the most common identified cause of cellulitis: a systematic review. Epidemiol Infect 138(3):313–317.

14. Jain, Seema Self, Wesley H. Wunderink, Richard G. Fakhran, Sherene Balk, Robert Bramley, Anna M. Reed, Carrie Grijalva, Carlos G. Anderson, Evan J. Courtney, Mark Chappell, James D. Qi, Chao (2015) Community-Acquired Pneumonia Requiring Hospitalization among U.S. Adults. N Engl J Med 373:415–427.

15. Wubbel L, et al. (1999) Etiology and treatment of community-acquired pneumonia in ambulatory children. Pediatr Infect Dis J 18(2):98–104.

16. Gwaltney JM Jr, Scheld WM, Sande MA, Sydnor A (1992) The microbial etiology and antimicrobial therapy of adults with acute community-acquired sinusitis: a fifteen-year experience at the University of Virginia and review of other selected studies. J Allergy Clin Immunol 90(3 Pt 2):457–61; discussion 462.

17. Brook I, Thompson DH, Frazier EH (1994) Microbiology and management of chronic maxillary sinusitis. Arch Otolaryngol Head Neck Surg 120(12): 1317–1320.

18. Celin SE, Bluestone CD, Stephenson J, Yilmaz HM, Collins JJ (1991) Bacteriology of acute otitis media in adults. JAMA 266(16):2249–2252.

19. Bluestone CD, Stephenson JS, Martin LM (1992) Ten-year review of otitis media pathogens. Pediatr Infect Dis J 11(Supplement):S7–11.

20. Edlin RS, Shapiro DJ, Hersh AL, Copp HL (2013) Antibiotic Resistance Patterns of Outpatient Pediatric Urinary Tract Infections. J Urol 190(1):222–227.

21. Gupta K, Scholes D, Stamm WE (1999) Increasing prevalence of antimicrobial resistance among uropathogens causing acute uncomplicated cystitis in women. JAMA 281(8):736–738.

